# Data-driven integration of hippocampal CA1 synapse physiology in silico

**DOI:** 10.1101/716480

**Authors:** András Ecker, Armando Romani, Sára Sáray, Szabolcs Káli, Michele Migliore, Audrey Mercer, Henry Markram, Eilif Muller, Srikanth Ramaswamy

**Affiliations:** Blue Brain Project, École polytechnique fédérale de Lausanne, Campus Biotech, Geneva, Switzerland; Institute of Experimental Medicine, Hungarian Academy of Sciences, Budapest, Hungary; Pázmány Péter Catholic University, Faculty of Information Technology and Bionics, Budapest, Hungary; Institute of Biophysics, National Research Council, Palermo, Italy; UCL School of Pharmacy, University College London, London, United Kingdom

**Keywords:** hippocampus, data integration, in silico modeling, CA1, synapse

## Abstract

The anatomy and physiology of synaptic connections in rodent hippocampal CA1 have been exhaustively characterized in recent decades. Yet, the resulting knowledge remains disparate and difficult to reconcile. Here, we present a data-driven approach to integrate the current state-of-the-art knowledge on the synaptic anatomy and physiology of rodent hippocampal CA1, including axo-dendritic innervation patterns, number of synapses per connection, quantal conductances, neurotransmitter release probability, and short-term plasticity into a single coherent resource. First, we undertook an extensive literature review of paired-recordings of hippocampal neurons and compiled experimental data on their synaptic anatomy and physiology. The data collected in this manner is sparse and inhomogeneous due to the diversity of experimental techniques used by different labs, which necessitates the need for an integrative framework to unify these data. To this end, we extended a previously developed workflow for the neocortex to constrain a unifying *in silico* reconstruction of the synaptic physiology of CA1 connections. Our work identifies gaps in the existing knowledge and provides a complementary resource towards a more complete quantification of synaptic anatomy and physiology in the rodent hippocampal CA1 region.

## 1 Introduction

The hippocampal CA1 region is probably the most studied region of the mammalian brain and is thought to play a pivotal role in learning and memory (Bliss and Collingridge, 2013; Buzsáki, 1989). Neuronal microcircuits in the hippocampal CA1 region process and store information through a myriad of cell-type-specific synaptic connections. Previous studies have identified that hippocampal cell-types are connected through multiple synapses, which are positioned across distinct axo-dendritic domains with a diversity of short- and long-term dynamics, as well as synaptic strengths. Despite the wealth of data, we lack an integrative framework to reconcile the diversity of synaptic physiology, and therefore, identify knowledge gaps. There have been several recent attempts to integrate knowledge about the hippocampal CA1 (Bezaire and Soltesz, 2013; Wheeler et al., 2015), however, they were not focused on the dynamics of synaptic transmission. Recent attempts have extended the utility of the online resource hippocampome.org towards synaptic electrophysiology as well (Moradi and Ascoli, 2019). However, in the continuing spirit of hippocampome.org, the study is primarily a text mining-based collection of papers and parameters, which does not integrate these data into a unifying framework. As a way forward, we extended a previously developed workflow to integrate disparate data on the physiology of synaptic transmission in hippocampal CA1, identified and extrapolated organizing principles to predict knowledge gaps (Markram et al., 2015). We accounted for the dynamic and probabilistic nature of synaptic transmission by fitting experimental traces using a stochastic generalization of the Tsodyks-Markram short-term plasticity model (Tsodyks and Markram, 1997; Markram et al., 1998; Fuhrmann et al., 2002). After validating the number and location of synapses, parameterizing the release probability and reversal potentials, as well as depression, facilitation, and synaptic conductance rise and decay time constant for various hippocampal connection types, we corrected for space clamp artefacts in silico by tuning synaptic conductance to match the in vitro PSP (postsynaptic potential) amplitudes. We also considered temperature and extracellular calcium concentration ([*Ca*^2+^]_*o*_) differences, which were adjusted using Q10 and Hill scaling factors, respectively. The resulting models for a subset of hippocampal connection types were applied predictively to the remaining uncharacterized connection types by clustering them into nine groups based on synapse types and neuronal biomarkers and applying the known parameters within each group. Curated and predicted parameters presented here should serve as a resource to researchers aiming to model hippocampal synapses at any level, while the detailed methodology intends to give a guideline to utilize such a framework to integrate data from other brain regions or species.

## 2 Methods

### 2.1 Circuit building and synapse anatomy

A detailed model of the rat hippocampal CA1 area was built using the pipeline of Markram et al. (2015). Circuit building and rigorous validation will be detailed in a following article. In brief, single cell models with detailed morphologies including axonal reconstructions from Migliore et al. (2018) were populated in an atlas-based volume corresponding to the dimensions of the hippocampal CA1 region. Structural appositions between axons and dendrites were detected based on touch distance criteria and were later pruned to match experimentally reported bouton density, number of synapses per connections and connection probability using an algorithm to yield the functional connectome (Reimann et al., 2015). In this manner, the number and location of synapses for each connection were constrained in a data-driven manner. Connected cell-types were sampled from this circuit based on their inter-somatic distance.

### 2.2 Dendritic features of single cell models

Detailed biophysical models of pyramidal cells (PCs) and interneurons of the CA1 region from Migliore et al. (2018) were re-optimized and used in the present study. All models were constrained with active dendritic conductances but were optimized using only somatic features (Migliore et al., 2018). While the somatic responses to various step-current injections were correct, the dendrites of the single cell models turned out to be too excitable (single synaptic input leading to spikelets and somatic spikes). For this reason, single cell models were re-optimized with slightly reduced range for dendritic sodium channel density. PSP propagation and attenuation along the dendritic branches is a key feature for our synaptic conductance calibration, thus it was validated against experimental data using the HippoUnit framework (unpublished). To this end, excitatory postsynaptic current (EPSC) like currents were injected into the apical trunk of PCs with varying distance from the soma and PSPs were simultaneously measured at the local site of the injection and in the soma.

### 2.3 Correcting for calcium ion concentration, temperature and liquid junction potential

Published parameters from different sources were corrected for differences arising from distinct experimental protocols. This included corrections for extracellular calcium levels different from 2 mM, temperatures different from 34 °Cand liquid junction potential (LJP) in the case of whole-cell recordings using patch pipettes. The correction for [*Ca*^2+^]_*o*_ was done by scaling the *U*_*SE*_ parameter of the synapses (see below), using the Hill isotherm with *n* = 4 (Hill, 1910):

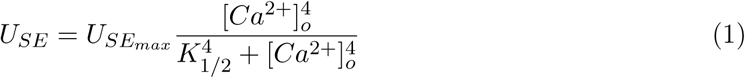

where *U*_*SE*_ is the absolute release probability and 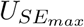 and *K*_1/2_ are free parameters. *K*_1/2_ values were taken from Rozov et al. (2001), 2.79 (mM) for steep and 1.09 (mM) for shallow calcium dependence and were shown to generalize well for other characterized pathways of the neocortex (see Supplementary Figure S11 in Markram et al. (2015)). In the absence of hippocampus specific data, we followed the approach of Markram et al. (2015) and assumed a steep dependence in PC to PC and PC to distal dendrite targeting inhibitory (O-LM) cells, and a shallow dependence between PC to proximal targeting cells (PV+BC (basket cell), CCK+BC, AAC). For experimentally uncharacterized pathways an intermediate calcium dependence was used, as the average of the steep and shallow ones. Temperature correction of kinetic parameters such as rise and decay time constants were realized by multiplying them with Q10 scaling factors:

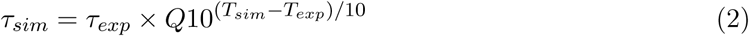

where *τ* is the time constant, Q10 is an empirically determined, receptor-specific parameter, *T*_*sim*_ = 34°C is the temperature used in the simulations, while *T*_*exp*_ is the temperature of the experiment. Holding potentials were corrected by the theoretical LJP (Neher, 1992). These potentials arise from the differences in solutions in the pipette and bath and are in 2-12 mV range for the standard solutions. Theoretical LJPs were calculated from the reported pipette and bath solutions with the Clampex 11 software.

### 2.4 Short-term plasticity model fitting

Short-term plasticity (STP) of synapse dynamics was fit by the Tsodyks-Markram (TM) model (Tsodyks and Markram, 1997). The model assumes that all synaptic connections have a finite amount of resources. Each presynaptic action potential utilizes a certain fraction of available resource (*R*) with a release probability (*U*), which then recovers. Over the years, the model has been refined and enriched to capture for example short-term facilitation and multi-vesicular release (MVR) (Markram et al., 1998; Loebel et al., 2009). The differential equations are as follows (see Supplementary Methods for comparison of different versions of the TM model):

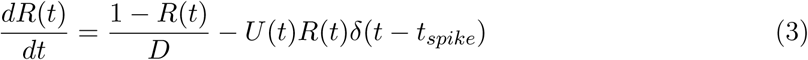

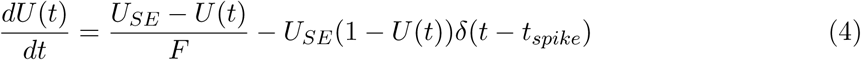

where *D*, and *F* and are depression and facilitation recovery time constants respectively, *U*_*SE*_ is the absolute release probability also known as the release probability in the absence of facilitation. *δ*(*t*) is the Dirac delta function and *t*_*spike*_ indicates the timing of a presynaptic spike. Each action potential in a train elicits an *A*_*SE*_*U* (*t*_*spike*_)*R*(*t*_*spike*_) amplitude PSC, where *A*_*SE*_ is the absolute synaptic efficacy. *R* = 1 and *U* = *U*_*SE*_ are assumed before the first spike. *U*_*SE*_, *D, F* and *A*_*SE*_ free parameters of the model were fit to experimentally recorded PSCs in Kohus et al. (2016) using a multiobjective genetic algorithm with BluePyOpt (Van Geit et al., 2016). Different frequency stimulations (10, 20 and 40 Hz) were fit together for better generalization. To correctly compare the coefficient of variation (CV, std/mean) of first PSC amplitudes, measurement noise was added to the simulated traces (Barros-Zulaica et al., 2019). To this end, noise parameters of *in vitro* traces were fitted and averaged for every different connection types and then stochastic noise generated with these extracted parameters was added to the corresponding *in silico* traces. Noise was described as an Ornstein-Uhlenbeck (OU) process. The OU process is a stationary Gauss-Markov process, which describes the velocity of the movement of a Brownian particle and is used in physics to describe noise relaxation (Bibbona et al., 2008). Mathematically it can be described with the following iterative equation:

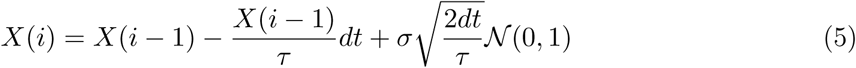

where *dt* is the time step of the signal, *τ* is the time constant fit to the exponential decay of the signal’s autocorrelation function, *σ* is the standard deviation of the signal and 𝒩 (0, 1) is a draw from the normal distribution.

### 2.5 In silico synapse model

The synapse model used in the simulations is based on the classical quantal model (Del Castillo and Katz, 1954), in which a synaptic connection is assumed to be composed of *N* independent release sites, each of which has a probability of release, *p* (function of *U*_*SE*_, *D, F*), and contributes a quanta *q* (function of the conductance *g*(*t*)) to the postsynaptic response (Ramaswamy et al., 2012, 2015; Markram et al., 2015; Chindemi, 2018). Conductances were modeled with double exponential kinetics:

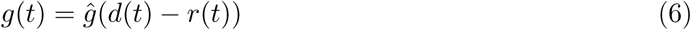

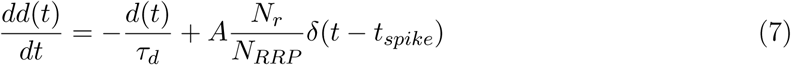

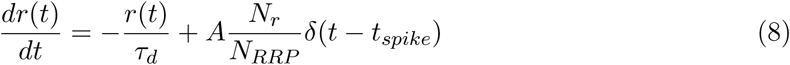

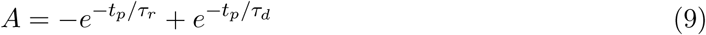

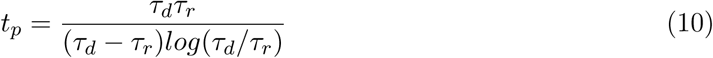

where *ĝ* (nS) is the peak conductance, *d* is the decaying component with time constant *τ*_*d*_ (ms) and *r* is the rising component with time constant *τ*_*r*_ (ms). Rise time constants are set to 0.2 ms for all pathways following Markram et al. (2015). Synapses were normalized (with A normalization constant) such as they reach peak conductance at time to peak *t*_*p*_ (ms). *N*_*r*_ is the number of released vesicles. Vesicle release dynamics was governed by a hybrid stochastic STP model (Fuhrmann et al., 2002). The model releases a single vesicle with probability *U* (*t*) (see TM model above) which then recovers. Vesicle recovery is an explicit process, meaning that compared to the canonical TM model, only fully recovered vesicles can be released. To this end, synaptic vesicles were implemented as 2-state (effective and recovering) Markov processes, in which staying in the recovered state at time *t* was described as a survival process, with time constant *D* (Chindemi, 2018):

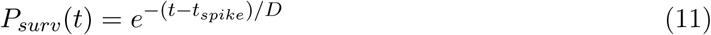

The above-described model converges to the canonical TM model in the limit (number of trials → ∞). MVR was implemented as *N* independent vesicles being released with the same probability *U* (*t*). *N*_*RRP*_ is the size of the readily releasable pool of vesicles and normalizing with it can be seen as scaling down the quantal size *q* of the quantal model in case of MVR, to keep the same mean PSP amplitudes, while changing only the variance (Barros-Zulaica et al., 2019). *N*_*RRP*_ was tuned to match the CVs of first PSC amplitudes from Kohus et al. (2016). Due to the lack of available raw data with STP protocol (and electron microscopy confirmation of the number of functional release sites) for most connections, the assumption of MVR (Conti and Lisman, 2003; Christie and Jahr, 2006) with *N*_*RRP*_ = 2 vesicles at each excitatory to excitatory terminal was used in this study, while all remaining non-tuned pathways were assumed to release single vesicles. See eg. Biro et al. (2005) and Gulyás et al. (1993) suggesting uni-vesicular release (UVR) for certain PC to interneuron connections. AMPAr and GABAr synaptic currents are then computed as:

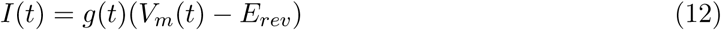

where *V*_*m*_ (mV) is the membrane potential and *E*_*rev*_ (mV) is the reversal potential of the given synapse. NMDAr currents depend also on *Mg*^2+^ block:

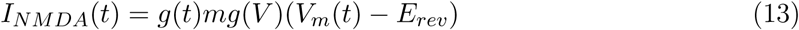

where *mg*(*V*) is the LJP corrected Jahr-Stevens nonlinearity (Jahr and Stevens, 1990):

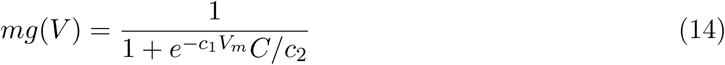

where *C* is the extracellular magnesium concentration and *c*_1_ = 0.062 (1/mV) and *c*_2_ = 2.62 (mM) are constants. NMDAr rise and decay time constants are Q10 corrected (Hestrin et al., 1990; Korinek et al., 2010) values from Andrasfalvy and Magee (2001): *τ*_*r*_ = 3.9 ms, *τ*_*d*_ = 148.5 ms. Peak NMDAr conductance *ĝ*_*NMDA*_ (nS) is calculated from the AMPAr one by multiplying it with NMDA/AMPA peak conductance ratio. NMDA/AMPA peak conductance ratio = 1.22 was taken from Groc et al. (2002); Myme et al. (2003). Synaptic currents are individually delayed based on axonal path length and conduction velocity of 300 *µ*m/ms (Stuart et al., 1997) and an additional 0.1 ms delay of neurotransmitter release (Ramaswamy et al., 2012).

### 2.6 Peak conductance tuning via in silico paired recordings

Paired recordings were replicated in silico as follows: Firstly, pairs were selected from the circuit based on distance criteria used by experimentalist (100 *µ*m cube for cells in the same layer and 200 *µ*m cube for cell pairs from different layers). Secondly, postsynaptic cells were current-clamped to match the LJP-corrected holding potential specified in the experiments. It is important to note, that in the case of pyramidal cells sodium channels were blocked (*in silico* TTX application) when clamping above −60 mV to avoid spontaneous firing of the cell models (see Figure 5 in Migliore et al. (2018)). Thirdly, a spike from the presynaptic cell was triggered, which stimulated all the synapses of the connection and resulted in a somatic PSP of the postsynaptic neuron. This exercise was run for 50 pairs with 35 repetitions for each. Lastly, mean PSP amplitude was compared to the experimentally reported one and peak conductance value was adjusted respectively using the formula:

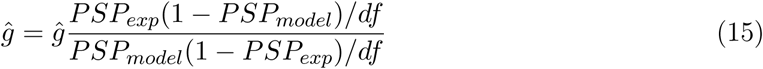

where *PSP*_*exp*_ (mV) and *PSP*_*model*_(mV) are the experimental and modeled PSPs amplitudes respectively and *df* = |*E*_*rev*_ − *V*_*hold*_| (mV) is the driving force. *E*_*rev*_ = 0 mV was used for excitatory connections, while *E*_*rev*_ = −80 mV for the inhibitory ones. All simulations were run using the NEURON simulator as a core engine (Hines and Carnevale, 1997) with the Blue Brain Project’s collection of .hoc and NMODL (Hines and Carnevale, 2000) templates for parallel execution on supercomputers (Hines et al., 2008a,b).

### 2.7 Statistical analysis

R values for validating matching experimental and model values are Pearson correlations. Data are presented as mean±std to yield comparable values to the experimental ones.

## 3 Results

The unifying workflow used to integrate synaptic data about the hippocampal CA1 is presented in Figure 1 and results from our literature review, parameter fitting and modeling will be detailed step-by-step in the following sections.

**Figure 1.**
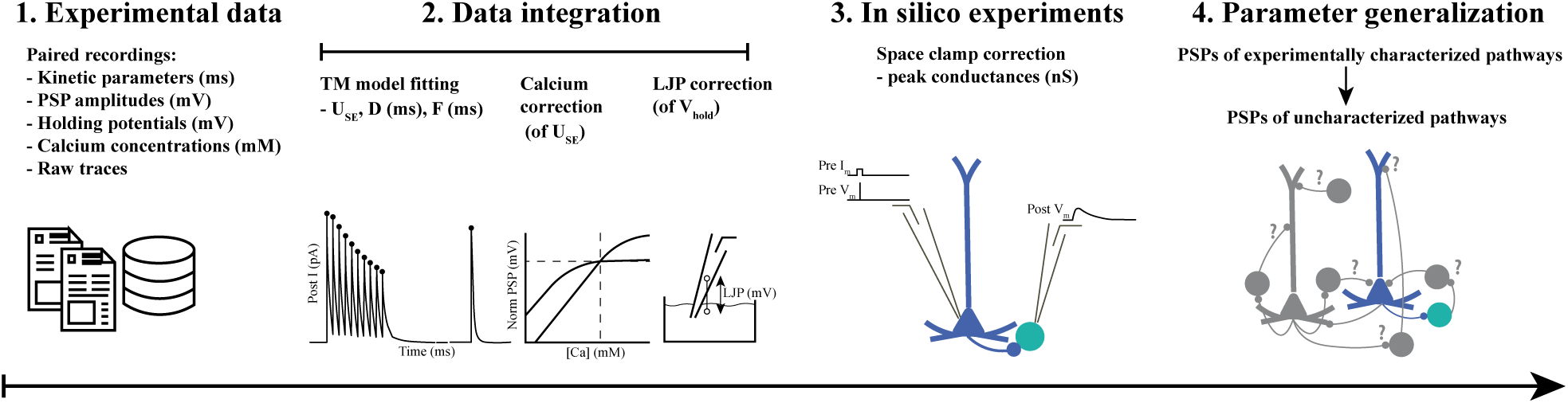
*In silico* data integration pipeline. **1:** More than a hundred publications were used to compile data on various parameters of connected neurons in rat CA1. **2:** Parameters were integrated into a common framework and experimental paradigm, including temperature, [*Ca*^2+^]_*o*_ and LJP corrections. TM models of STP were fit to publicly available raw traces. **3:** *In silico* paired recordings were run to correctly estimate the unitary peak conductance of connections with experimentally characterized PSP amplitudes. **4:** Parameters were averaged within classes and used predictively to describe experimentally uncharacterized pathways.

### 3.1 Literature curation

Firstly, we undertook an extensive literature review of paired recording experiments, and compiled data on the various parameters (see Supplementary Table S1 for voltage clamp, and Supplementary Table S2 for current clamp recordings from rat hippocampal CA1). The data collected in this manner is sparse and inhomogeneous, due to the diversity of experimental conditions used by different labs and were corrected for various aspects. [*Ca*^2+^]_*o*_ is known to affect release probability and additional Hill scaling had to be considered when parametrizing STP profiles (see Methods). Rise and decay time constants of synaptic currents are influenced by temperature differences but can be corrected with Q10 factors (see Methods). For electrophysiological recordings patch pipettes are becoming the standard practice over sharp electrodes nowadays, however, care should be taken when using absolute potentials reported from publications using whole-cell patch-clamp recordings (see Methods).

### 3.2 Validation of synapse anatomy and dendritic attenuation

Before we ran any simulations with synapses using the extracted parameters, we verified that the anatomy of synapses (Figure 2) such as the number of synapses per connection and targeting profile, as well as basic electrophysical properties of the cell models match experimental data. Cell pairs used in the simulations were pulled out from a data-driven reconstruction of the rat CA1 region, built with the pipeline presented in Markram et al. (2015). Number of synapses per connection for experimentally characterized pathways (Ali, 2011; Biro et al., 2005; Buhl et al., 1994a,b; Deuchars and Thomson, 1996; Földy et al., 2010; Maccaferri et al., 2000; Sik et al., 1995; Vida et al., 1998) (*r* = 0.98, Figure 2 b and Supplementary Table S3) along with targeting profile (Figure 2 a) was verified for this work. PSP attenuation in the active dendrites of PCs (Migliore et al., 2018) is also in line with the experimentally reported curves (Magee and Cook, 2000) (Supplementary Figure S1).

**Figure 2.**
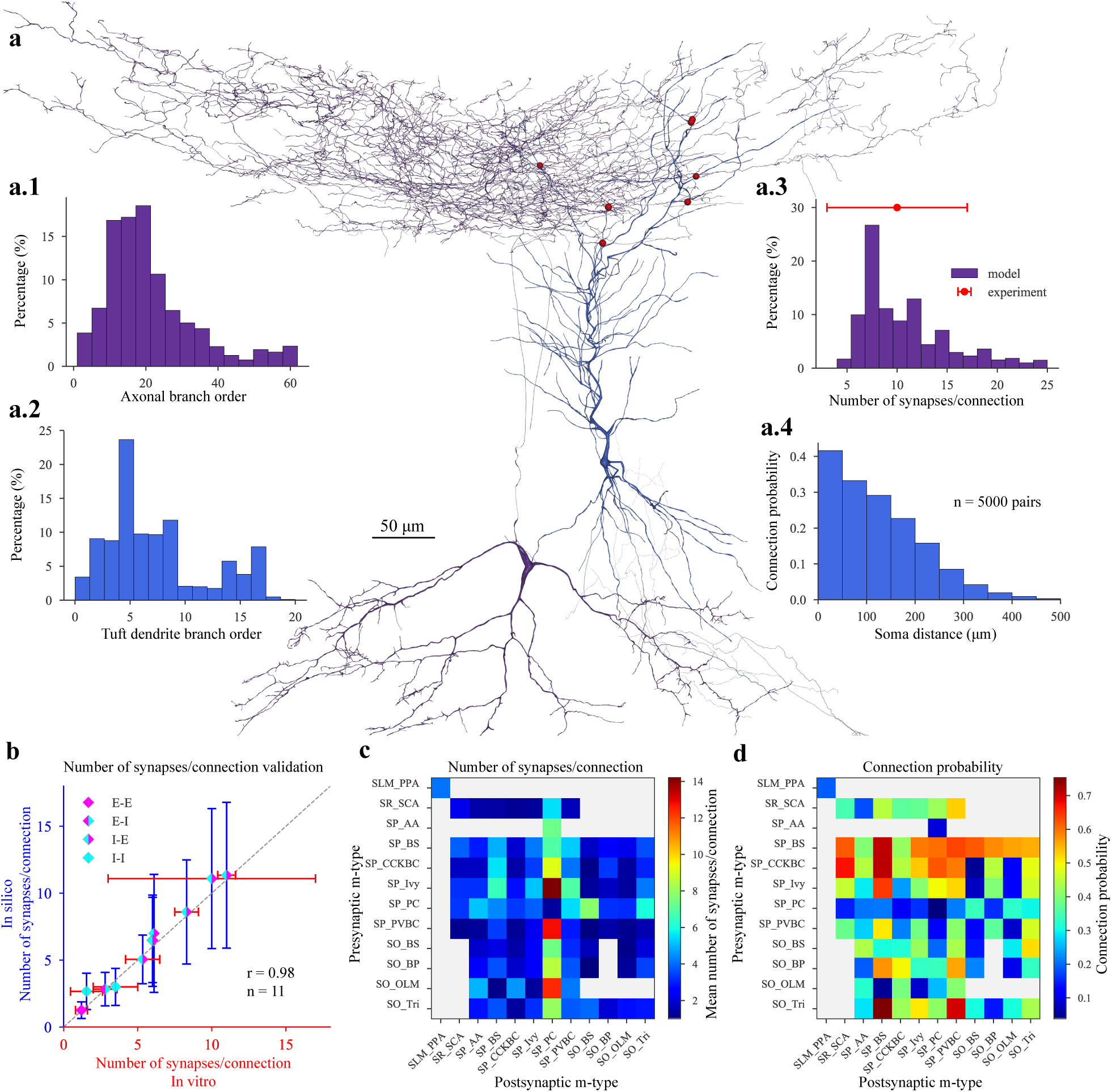
*In silico* synapse anatomy. **a:** A representative *in silico* O-LM (purple) to PC (blue) pair, with synapses visualized in red. 3D morphologies were reconstructed with the Neurolucida software (Migliore et al., 2018) by the members of Alex Thomson’s lab at UCL. **a.1:** Branch order distribution (n=5000 connections) of the presynaptic (O-LM) axons. **a.2:** Branch order distribution of the postsynaptic (PC) tuft dendrites. **a.3:** Distribution of the number of synapses per connection of the *in silico* O-LM to PC pathway. *In vitro* experimental data is indicated in red. **a.4:** Distance dependent connection probability of the *in silico* O-LM to PC pathway. **b:** Validation of the number of synapses per connection against experimental data. (E: excitatory, I: inhibitory, eg.: I-E: inhibitory to excitatory pathways.) **c:** Predicted mean number of synapses per connections (within 200 *µm* intersomatic distance) for all pathways in the CA1 network model. Layer abbreviations: SR: stratum radiatum, SP: stratum pyramidale, SO: stratum oriens. M-type (morphological type) abbreviations: AA: axo-axonic cell, BP: back-projecting cell, BS: bistratified cell, CCKBC: CCK+ basket cell, Ivy: ivy cell, OLM: oriens-lacunosum moleculare cell, PC: pyramidal cell, PVBC: PV+ basket cell, PPA: performant path-associated cell, SCA: Schaffer collateral-associated cell, Tri: trilaminar cell. **d:** Predicted mean connection probability (within 200 *µm* intersomatic distance) for all pathways in the CA1 network model. M-type abbreviations are as in **c**.

### 3.3 Short-term plasticity of synapses

Transmission properties of hippocampal CA1 neurons were demonstrated to express a wide range of STP profiles in response to presynaptic trains of action potentials at different frequencies (Ali et al., 1998, 1999; Ali and Thomson, 1998; Éltes et al., 2017; Kohus et al., 2016; Losonczy et al., 2002; Pouille and Scanziani, 2004). However, to our best knowledge, only Losonczy et al. (2002) and Kohus et al. (2016) reported TM model parameters for CA1 pathways and used additional recovery spike after the spike train, which is crucial to distinguish pseudo-linear profiles from purely facilitating or depressing ones. Published STP parameters from Losonczy et al. (2002) were used for PC to basket cell pathways, after refitting a subset of their data and confirming the similarity between our resulting *U*_*SE*_, *D, F* values. Kohus et al. (2016) also took the effort to make their raw traces publicly available, thus despite all the differences from our standard approach (current-clamp recordings from rat at [*Ca*^2+^]_*o*_ = 2 mM) we used their traces to fit the parameters (see Methods) of the TM model (Tsodyks and Markram, 1997; Markram et al., 1998). We rigorously validated that our event-based amplitude fitting is equivalent to the equations previously presented in the literature (Markram et al., 1998; Maass and Markram, 2002) (see Supplementary Methods). Fitted parameters matched well the ones fitted in the original article (Kohus et al., 2016), despite the slight differences in the TM model used, and the CVs of the first PSC amplitudes, which were not used during the fitting (see Methods) were also close to experimental ones (*r* = 0.8, Figure 3 b, Supplementary Table S4). (CCK+ dendrite targeting interneurons were used as Schaffer collateral-associated cells.) For PV+BC to PC and PV+BC we had to introduce MVR (see Methods) (with *N*_*RRP*_ = 6 vesicles) to match the CVs of the measured PSCs (Figure 3 b). On the other hand, *in silico* PV+BC to AA PSCs had lower CVs with UVR than the *in vitro* ones, which could not be corrected. (MVR can reduce the variance, but not increase). Biró et al. (2006) have shown in an elegant study, that while CCK+BC to PC connections in CA3 are characterized by MVR (with *N*_*RRP*_ = 5 − 7 vesicles) under experimentally imposed high release probability conditions (high extracellular Ca/Mg ratio), at physiologically relevant [*Ca*^2+^]_*o*_ UVR is more prevalent. In our simulations, the CV of the *in silico* CCK+BC to PC PSCs matched well the *in vitro* ones, recorded under physiological conditions using UVR, in good agreement with the Biró et al. (2006) study. For the remaining pathways *U*_*SE*_, *D, F* values from the analogous pathways of the somatosensory cortex (Markram et al., 2015) were used since parameters of the comparable connections matched well (perisomatic inhibitory to PC, inhibitory to inhibitory) and that is the most comprehensive dataset available to date. Based on the literature and our model-fitting we identified several rules to characterize and group STP profiles. The characterization of all pathways result as follows (Table 1, Figure 4): PC to O-LM cells (Ali and Thomson, 1998; Biro et al., 2005; Losonczy et al., 2002; Pouille and Scanziani, 2004) and other interneurons in stratum oriens (Éltes et al., 2017) E1 (excitatory facilitating). PC to PC (Deuchars and Thomson, 1996), PC to all SOM-interneurons (Ali et al., 1998; Losonczy et al., 2002; Pouille and Scanziani, 2004) E2 (excitatory depressing). CCK+ interneurons to CCK+ interneurons (Ali, 2007, 2011; Kohus et al., 2016) I1 (inhibitory facilitating), PV+ and SOM+ interneurons to PC (Ali et al., 1998, 1999; Bartos et al., 2002; Buhl et al., 1995; Daw et al., 2009; Kohus et al., 2016; Maccaferri et al., 2000; Pawelzik et al., 2002) as well as interneurons to interneurons (except the CCK+ ones) (Bartos et al., 2002; Daw et al., 2009; Elfant et al., 2008; Karayannis et al., 2010; Kohus et al., 2016; Price et al., 2005) I2 (inhibitory depressing). CCK+ and NOS+ (only Ivy cells, since we lack NGF morphologies) to PC (Fuentealba et al., 2008; Kohus et al., 2016; Price et al., 2008) I3 (inhibitory pseudo linear). It is important to note here that these profiles are valid in juvenile animals at [*Ca*^2+^]_*o*_ = 2 mM, but in some cases, release probability scales drastically with [*Ca*^2+^]_*o*_ and the STP profiles can change as well. For example, at an *in vivo* like calcium level (1.1-1.3 mM) the PC to PC pathway can show an E3 (excitatory pseudo-linear) characteristic with amplitudes having a lower mean and higher trial-by-trial variability and more failures compared to the *in vitro* (2 mM) depressing E2 profile (Supplementary Figure 2 b). As a function of [*Ca*^2+^]_*o*_ *U*_*SE*_ values (absolute release probability parameter of the TM model) are scaled by Hill isotherm (see Methods) parametrized with cortical data of PSP amplitude changes (Supplementary Figure S11 in Markram et al. (2015)). Here we have shown that applying this scaling function on the absolute release probabilities indeed results in the same scaling profile of PSP amplitudes in the case of PC to PC connection (Supplementary Figure 2 a).

**Table 1:**
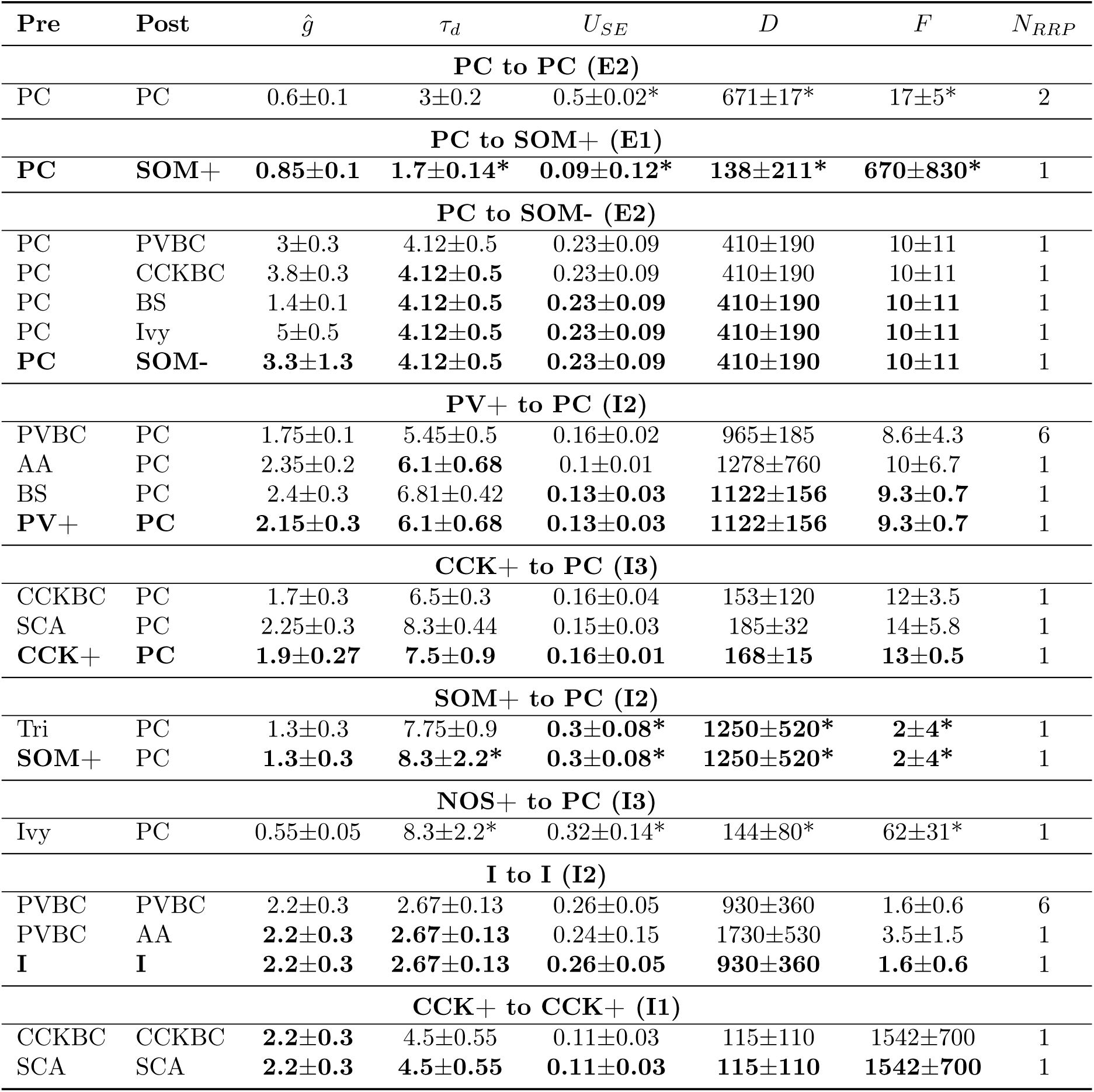
Parameters and generalization to 9 classes. Table with model synapse parameters either extracted from the literature (*τ*_*d*_ (ms)), fitted (*U*_*SE*_, *D* (ms), *F* (ms)), tuned (*ĝ*(nS)) or taken from the somatosensory cortex (marked with *) (Markram et al., 2015). Average class parameters are marked in bold and are used predictively for the remaining pathways belonging to the same class. Abbreviations are as in **Figure 2 b**

**Figure 3.**
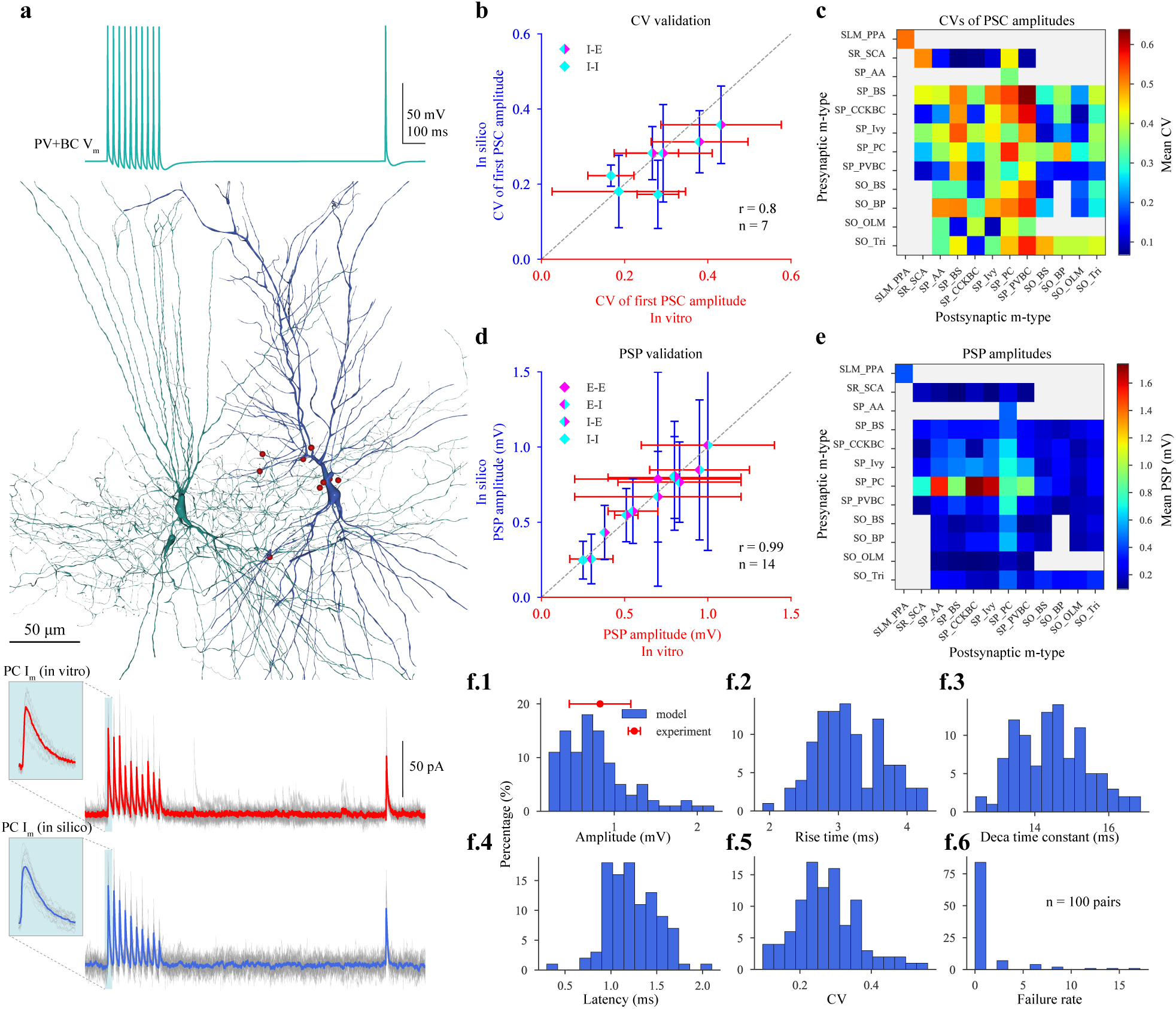
*In silico* synapse physiology. **a:** *In silico* paired recording experiment with the STP protocol used in Kohus et al. (2016). Presynaptic (PV+BC) voltage trace is shown on top. In silico PV+BC (green) to PC (blue) pair, with synapses visualized in red in the middle. 3D morphologies were reconstructed with the Neurolucida software (Migliore et al., 2018) by the members of Alex Thomson’s lab at UCL. Postsynaptic (PC) experimental traces recorded in vitro (in gray) and their mean in red, as well as model traces recorded in silico (in gray) and their mean in blue, are presented at the bottom panel. Insets show the variance of the first IPSCs. **b:** Validation of the CV of the first PSC amplitudes against experimental data. (E: excitatory, I: inhibitory, eg.: I-E: inhibitory to excitatory pathways.) **c:** Predicted CVs of first PSC amplitudes for all pathways in the CA1 network model after synapse parameter generalization. 20 pairs with 35 repetitions for every possible connection. Postsynaptic cells were held at −65 mV in *in silico* voltage-clamp mode. M-type abbreviations are as in **Figure 2 b. d:** Validation of the PSP amplitudes against experimental data. (E: excitatory, I: inhibitory, eg.: I-E: inhibitory to excitatory pathways.) **e:** Predicted PSP amplitudes for all pathways in the CA1 network model after synapse parameter generalization. 20 pairs with 35 repetitions for every possible connection. Postsynaptic cells were held at −65 mV in *in silico* current-clamp mode. M-type abbreviations are as in **Figure 2 b. f:** Properties of postsynaptic (PC) IPSPs. **f.1:** Distribution of *in silico* PSP amplitudes. *In vitro* experimental data is indicated in red. **f.2:** Distribution of *in silico* PSP 10-90% rise times. **f.3:** Distribution of *in silico* PSP decay time constants. **f.4:** Distribution of *in silico* PSP latencies. **f.5:** Distribution of the CVs of the first *in silico* PSP amplitudes. **f.6:** Distribution of *in silico* failure rates.

**Figure 4.**
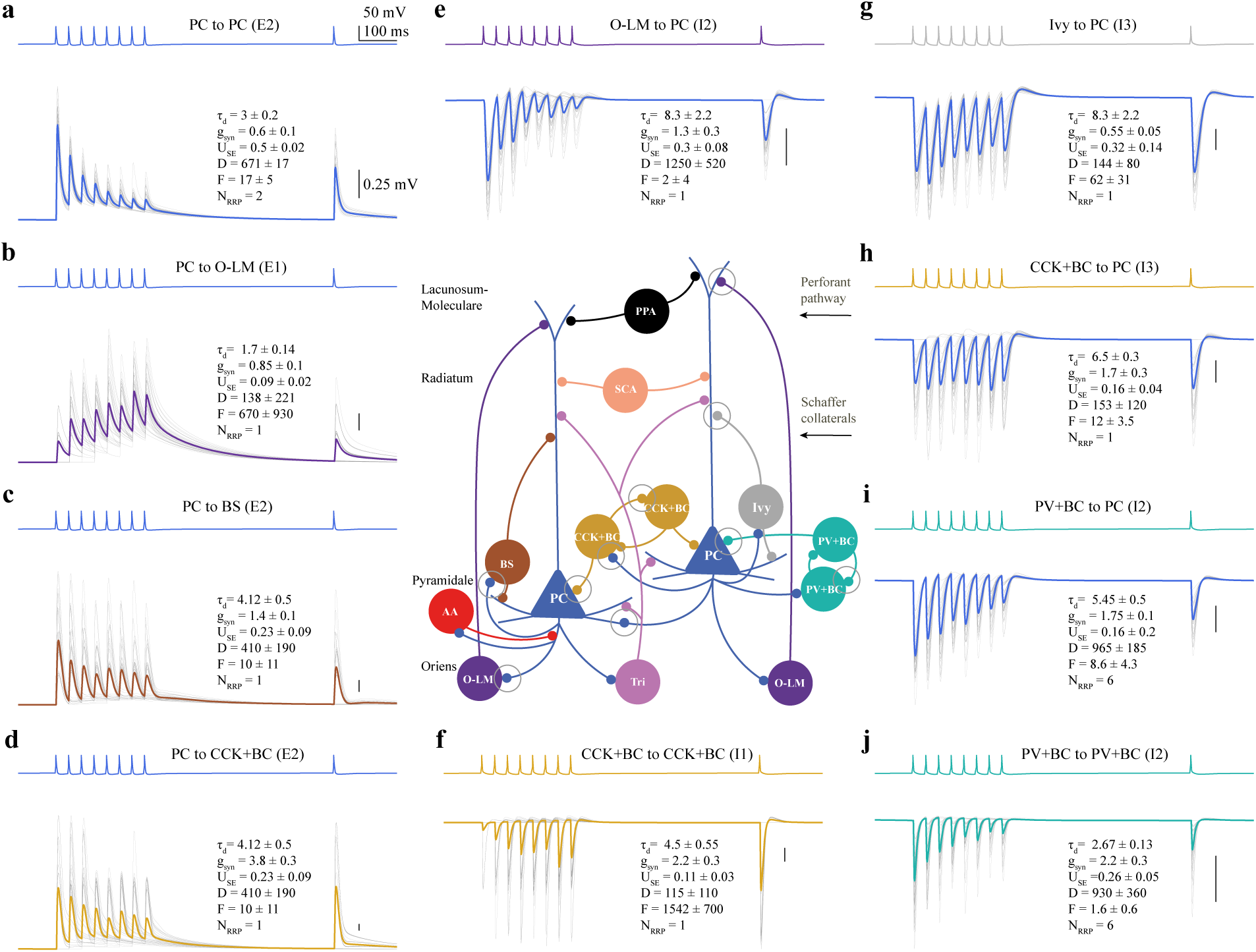
Summary of synapse diversity in the CA1 network model. Panels represent exemplar *in silico* pairs from the 9 generalized pathways (2 for PC to SOM-interneurons). Presynaptic voltage traces are shown on the top of the panels, while 35 repetitions (in gray) and their mean postsynaptic PSPs are presented on the bottom of the panels for each pathway. Postsynaptic cells were held at −65 mV in *in silico* current-clamp mode. **a:** PC to PC (E2). **b:** PC to O-LM cell (E1). **c:** PC to bistratified cell (E2). **d:** PC to CCK+BC (E2). **e:** O-LM cell to PC (I2). **f:** CCK+BC to CCK+ BC (I1). **g:** Ivy cell to PC (I3). **h:** CCK+BC to PC (I3). **i:** PV+BC to PC (I2). **j:** PV+BC to PV+BC (I2). Connectivity in the schematic CA1 microcircuit in the middle is simplified for visualization purpose (for example most of the interneuron to interneuron connections are missing). Simplified synapses of the pathways shown in the panels around are indicated with gray circles. M-type abbreviations are as in **Figure 2 b**.

### 3.4 Tuning of peak conductances to match PSP amplitudes

Peak conductances of single synapses cannot be measured routinely with today’s experimental techniques, thus are always somehow tuned to match a desired behavior in modeling studies. While it is appealing to calculate peak conductances from voltage-clamp recordings simply by dividing peak PSC amplitudes by the driving forces and plug them into a synapse model, it should not be done because of the space clamp artifacts (Bar-Yehuda and Korngreen, 2008; Spruston et al., 1993; Williams and Mitchell, 2008). Namely, if one voltage clamps the soma of a neuron, that will not necessarily mean that the dendritic compartments where most of the synapses arrive will have the same holding voltage (which cannot be compensated experimentally) and this can bias the driving force estimate. Furthermore, in the case of thin dendrites and strong synapses, the relation between the PSC amplitude and the peak conductance is rather sublinear (Gulyás et al., 2016). Using the same reasoning and access to connections measured in both voltage clamp and current clamp modes from the somatosensory cortex we have recently shown that the space clamp corrected *in silico* peak conductances are at least twice as big as their calculated counterparts (Markram et al., 2015). In the case of rat hippocampal CA1, we did not have the luxury of having both PSCs and PSPs from the same pair (See Supplementary Tables S1 and S2), thus just used all PSPs to tune the in silico peak conductances to match the *in vitro* PSPs (Ali et al., 1998; Ali and Thomson, 1998; Cobb et al., 1997; Deuchars and Thomson, 1996; Fuentealba et al., 2008; Pawelzik et al., 1999, 2002) (Figure 3 d, Table 1). In short, all other synapse parameters (anatomy, rise, and decay time constants, STP parameters, *N*_*RRP*_, NMDA/AMPA peak conductance ratio, reversal potential) were rigorously validated, a pair was selected from the digitally reconstructed circuit, the postsynaptic neuron was current clamped to the given holding voltage, a spike was delivered from the presynaptic neuron, which caused a PSP, measured in the soma. After repeating this for multiple pairs (*n* = 50) with many trials for each (*n* = 35) we scaled the peak conductance to match the reference mean PSP amplitude (see Methods). Next, we repeated the same protocol on a different set of randomly selected 50 pairs with the tuned peak conductance distributions as a validation of the reconstruction process itself (*r* = 0.99, Figure 3 d, Supplementary Table S5). As an external validation of the resulting peak conductances, we set to compare them to published single-channel conductance and receptor number estimates. We only found sufficient data in the case of excitatory connections to PCs. CA1 PCs receive most of their excitatory inputs from CA3 PCs by the Schaffer collaterals (Megías et al., 2001; Takács et al., 2012), whereas in the present article we only considered internal connections (eg. excitatory connections between CA1s) and no long-range projections. Thus, single-channel conductance and receptor number estimates from the Schaffer collateral synapses were assumed to generalize for the internal PC to PC connections (Table 2). Using non-stationary fluctuation analysis on EPSCs recorded in outside-out dendritic membrane patches, Spruston et al. (1995) estimated 10.2 pS AMPA and 43.5 pS NMDA single-channel conductances. Using these numbers, our tuned 0.6 ± 0.1 nS AMPA peak conductance (Table 1) is the net result of ∼59 AMPA and ∼18 NMDA receptors (with 1.33 NMDA/AMPA peak conductance ratio, see Methods). In their *in vitro* study, Spruston et al. (1995) estimated 58 − 70 AMPA and 5 − 30 NMDA receptors (Jonas et al., 1993), which align well with our *in silico* predictions. MPA receptor numbers were also estimated with a quantitative immunogold localization technique (Nusser et al., 1998), as well as by non-stationary fluctuation analysis on single-spine level following two-photon glutamate uncaging (Matsuzaki et al., 2001) and these numbers also parallel with our predictions. Taken together, these data serve as an independent validation of the tuned peak conductance of the most important, PC to PC pathway. Predicted average GABA conductance is 1.8 ± 0.6 nS, which corresponds to ∼90 GABA receptors, which is also in good agreement with general estimates for the central nervous system (Mody and Pearce, 2004).

**Table 2:**
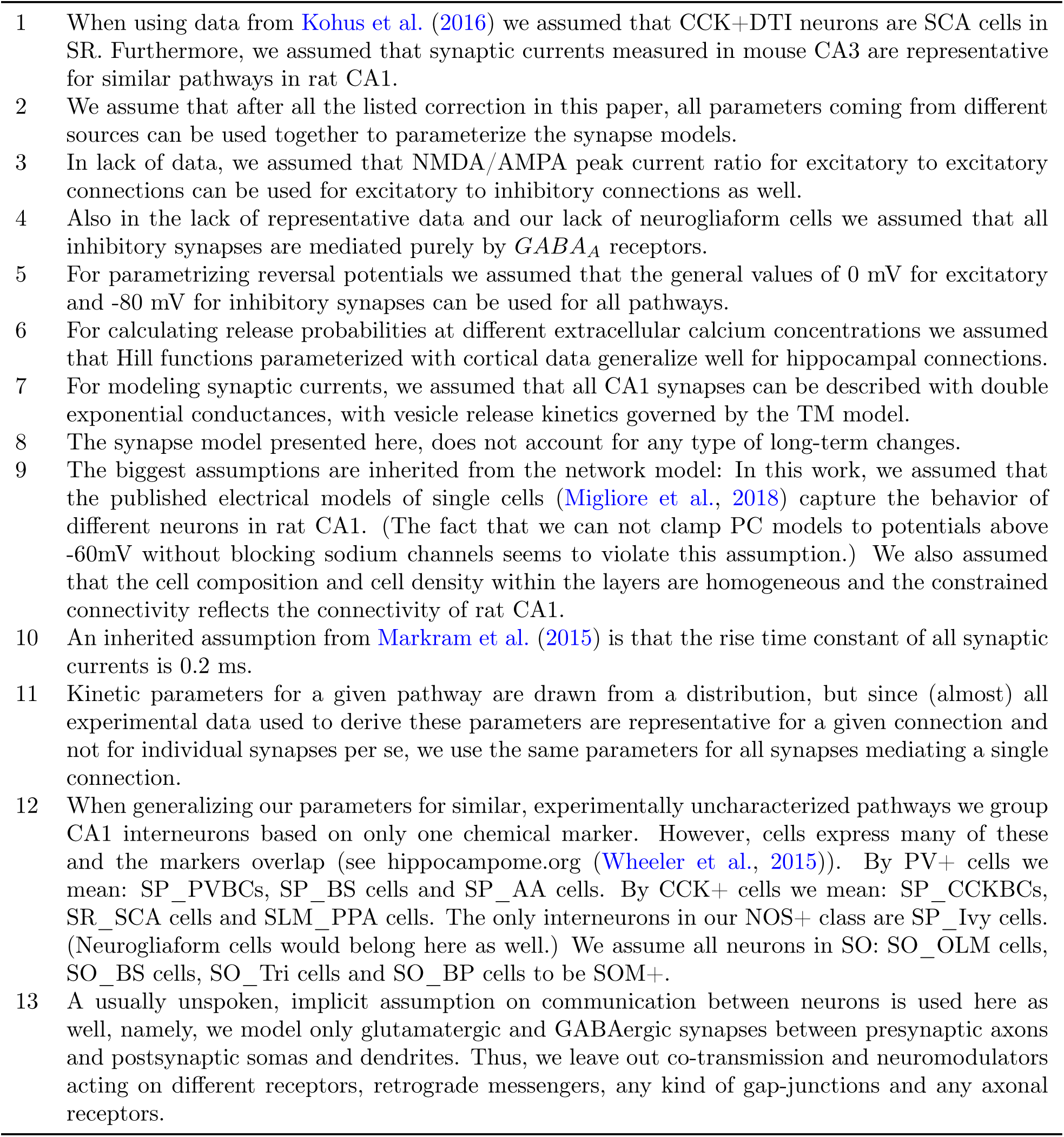
List of assumptions. All the assumptions that were made to arrive at model parameters from a sparse set of raw data and published values.

### 3.5 Parameter generalization

After integrating all the parameters (Table 1), obtaining values from somatosensory cortex to fill knowledge gaps when necessary (Table 1), and simulating paired recordings *in silico* we could extend predictions derived through this framework to other pathways (Figure 3 c, e). Synapse anatomy of the experimentally uncharacterized pathways was assumed to be correct and missing kinetic parameters were filled in with average values from the known ones, grouped by neurochemical markers, targeting and STP profiles and peak conductances (Table 1). This exercise resulted in 9 classes, covering all connection types in the CA1 region (Table 1, Figure 4). All the assumptions used in this study leading to the set of presented model parameters are listed in Table 2. Among other values, we predicted the first PSP amplitudes of all possible connections (Figure 3 e), given our cell models (Migliore et al., 2018) and connectivity. An exemplar rare case of more than one published value for a given synaptic property in the literature is the notion of “strong” connection between PCs and CA1 interneurons (Gulyás et al., 1993), which could not be used directly for tuning because the postsynaptic target was not clear, but was confirmed after generalization and *in silico* experiments with all possible postsynaptic interneurons (Figure 3 e). It is important to note, that due to the highly detailed nature of our digital, data-driven reconstruction process not only mean pathway values (Figure 3 c, e) but also detailed distributions can be predicted with the framework (Figure 3 f).

### 3.6 Synaptic strength

It is general practice among modelers using simplified models to represent synapses as single contacts between neurons and parameterize them with a single “weight”. It is important to note that in the detailed models presented here to concept of “weight” is a result of several features not just the peak conductance. This concept depends on the number and location of synapses (Figure 2), dendritic attenuation (Supplementary Figure S1), NMDA/AMPA conductance ratio and the interplay between release probability, number of vesicles and peak conductance. As an example, connections made by interneurons targeting perisomatic regions of PCs are mediated by multiple synapses with almost no attenuation, however have large peak conductances to compensate for the relatively low release probability (Table 1, Figure 4 h, i). They are characterized by low *U*_*SE*_ values (in our stochastic model the release probability almost always equals *U*_*SE*_ for depressing connections) and thus a high trial-to-trial variance. A notable exception from this high trial-to-trial variance is the PV+BC to PC pathway (Figure 3 a, f), which is more reliable (Figure 3 f.6) thanks to the MVR with 6 independent synaptic vesicles per a single synapse.

## 4 Discussion

Recent advances in high-performance computing have enabled biologically detailed, data-driven reconstructions and large-scale simulations of brain regions (Bezaire and Soltesz, 2013; Bezaire et al., 2016; Markram et al., 2015; Wheeler et al., 2015). In the present study, we have demonstrated that a data-driven workflow grounded in biological first-principles, which was used to digitally reconstruct a biologically detailed model of rat neocortical tissue, can be extended to model other brain regions such as the hippocampal CA1, to reconcile disparate cellular and synaptic data, and to predictively extrapolate the sparse set of parameters to synaptic connections that have not yet been characterized experimentally. It is known that [*Ca*^2+^]_*o*_ regulates the neurotransmitter release probability, and therefore, the amplitudes of PSPs. In this study we adapted the existing data-driven digital reconstruction workflow to reconcile differences in synaptic dynamics that were characterized at different levels of extracellular calcium. Therefore, we scaled the neurotransmitter release probabilities for all pathways that were characterized at 1.6-2 mM [*Ca*^2+^]_*o*_ (Kohus et al., 2016; Losonczy et al., 2002; Markram et al., 2015) before tuning peak conductances to match PSP amplitudes that were measured at 2.5 mM [*Ca*^2+^]_*o*_, which is more representative of baseline values for slice experiments (Ali et al., 1998; Ali and Thomson, 1998; Deuchars and Thomson, 1996; Fuentealba et al., 2008; Pawelzik et al., 1999, 2002).

In the continuing spirit of unifying hippocampal synaptic electrophysiology from published literature a recent complementary study leveraged text-mining techniques to extract the properties of synaptic connections in hippocampal CA1, including PSP amplitudes and peak conductances (Moradi and Ascoli, 2019). However, our approach to data integration from literature demonstrates that synaptic properties reported in the literature such as peak conductances should not be interpreted on face value but require further corrections to account for inadequate space-clamp errors, which could severely underestimate their value by two-three fold (Markram et al., 2015). The results we report, to the best of our knowledge, constitute a comprehensive resource for not only for the anatomy but also the kinetic and short-term dynamic physiological properties of the rat hippocampal CA1 region. Consolidation of the state of the literature not only facilitates building detailed models, but also highlights knowledge gaps and could help in prioritizing the identification of missing data on CA1 connections, such as PC to interneurons, and between interneurons, which are key building blocks of feedback inhibition. Indeed, the parameter set presented here should be considered a first draft, which will be systematically refined as and when new experimental data become available. By detailing all the integration steps in this study, we had two main objectives. First, we aimed to demonstrate that published parameters should not be taken at face value without rigorously checking their consistency within any modeling framework, and the necessity of being abreast of the state-of-the-art experimental techniques. Second, we attempted to emphasize the fact that a growing diversity of experimental standards combined with published literature that provides access to only processed data sets but not raw experimental traces could lead to an inconsistent picture of a fundamental mechanism such as synaptic transmission. The bottom-up modeling framework presented as a resource in this article could enable ways to integrate disparate data and provide a platform in catalyzing community-driven consensus on the synaptic organization of the hippocampal formation.

## Acknowledgements

We would like to thank Giuseppe Chindemi, Natali Barros-Zulaica, Rodrigo Perin and Zoltán Nusser for fruitful discussions as well as Werner Van Geit, Michael Gevaert, Arseniy Povolotsky, Cyrille Favreau and Marwan Abdellah for technical assistance. An initial version of this manuscript was submitted to bioRxiv.

## Author contribution

S.R., E.M. and A.R. conceptualized and supervised the study. S.K., M.M., A.M. and H.M. contributed to the supervision of the study. A.R. built the CA1 circuit with inputs from all authors. S.S. validated PSP attenuation. A.E. curated literature, performed simulations, analysis and created the figures with inputs from S.R. A.E. and S.R. wrote the manuscript with inputs from all authors.

## Supplementary Material

### Supplementary Methods

The Tsodyks-Markram model of short-term plasticity underwent many changes in the years twenty years. For a recent and consistent review see Hennig (2013). Furthermore, the equations are sometimes shown in the form of differential equations (Tsodyks and Markram, 1997; Tsodyks et al., 2000; Fuhrmann et al., 2002, 2004; Loebel et al., 2009; Hennig, 2013), while in other papers the iterative solution evaluated at spike arrivals is presented (Markram et al., 1998; Maass and Markram, 2002). The version used in this article follows the formalism presented in Hennig (2013):

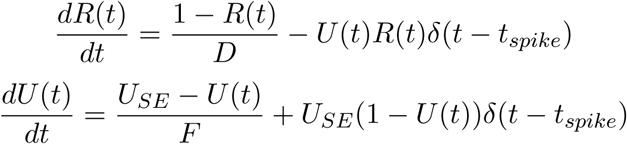

where *R*(*t*) is the fraction of available resources, *U* (*t*) is the release probability, *D*, and *F* are depression and facilitation time constants respectively. *U*_*SE*_ is the absolute release probability also known as the release probability in the absence of facilitation. *δ*(*t*) is the Dirac delta function and *t*_*spike*_ indicates the timing of a presynaptic spike. Each action potential in a train elicits an *A*_*SE*_*U* (*t*_*spike*_)*R*(*t*_*spike*_)amplitude PSC, where *A*_*SE*_ is the absolute synaptic efficacy and is linked to the *Nq* part of the quantal model, where *N* is the number of release sites and *q* is the quantal amplitude. *R* = 1, and *U* = *U*_*SE*_ are assumed before the first spike. In our simulations, we implement Fuhrmann et al. (2002) as the stochastic generalization of the model. The equation of the release probability is slightly different in that article and it reads as follows:

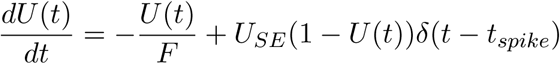

According to this equation *U* (*t*) decays to 0 (the wording of the articles suggest a decay to *“the baseline”*). To recover the definition of *U*_*SE*_ as the release probability in absence of spikes (or U as the constant release probability in the first Tsodyks and Markram (1997) paper concentrating only on depressing connections) the +*U*_*SE*_ (1 − *U* (*t*)) has to be evaluated before the release happens. On the other hand, the −*U* (*t*)*R*(*t*) jump in the equation of R still has to be evaluated after the event in order to be consistent with *R* being 1 in the absence of spikes. In this view *U* (*t*) is mostly zero and at spike arrivals, before release happens it jumps to *U*_*SE*_. From the biophysical point of view, this can be seen as a calcium-based model, where a quick calcium influx leads to release. On the other hand, in the Hennig (2013) version *U* (*t*) decays to its baseline *U*_*SE*_ value and the *U*_*SE*_ (1 −*U* (*t*)) jump happens after the release. When fitting the deterministic TM model to experimental data as well as simulating the stochastic version we use an event-based solution, meaning that the equations are only evaluated at spike times (as opposed to the ODE form).

For the Fuhrmann et al. (2002) version the iterative update is:

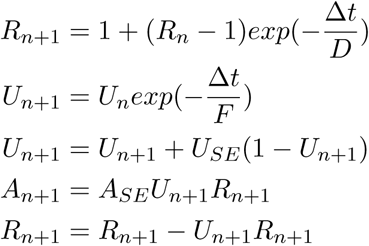

where Δ*t* is the the time between the (*n* + 1)th and *n*th spike and *A*_*n*_ is the *n*th amplitude. On the other hand, the Hennig (2013) version (used to fit models in Kohus et al. (2016)) is:

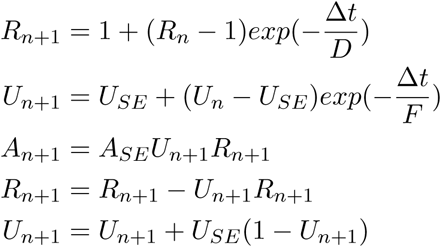

None of these forms are presented in the literature. Both Markram et al. (1998) and Maass and Markram (2002) put the jump terms into the decaying exponential part as follows:

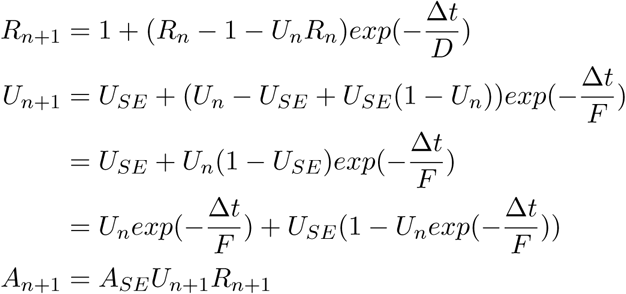

Using the initialization *R*_1_ = 1, *U*_1_ = *U*_*SE*_ and calculating the first two a mplitudes w ith a ll 3 versions (Fuhrmann et al. (2002), Hennig (2013) and Maass and Markram (2002)) one gets:

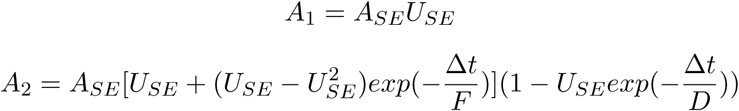

With simulations, it is also possible to show that all the other amplitudes in response to a spike train will be the same for all versions. Thus, the 3 event-based models presented above are equivalent even if it would be hard to confirm by algebra. We present the Hennig (2013) formalism in the article since we find it more intuitive that both Dirac deltas are evaluated at the same point (after the PSC amplitude is calculated) and is more in line with the wording of the papers, but emphasize that it is consistent with the other versions and the fits presented in Markram et al. (2015).

## Supplementary Figures

**Figure S1:**
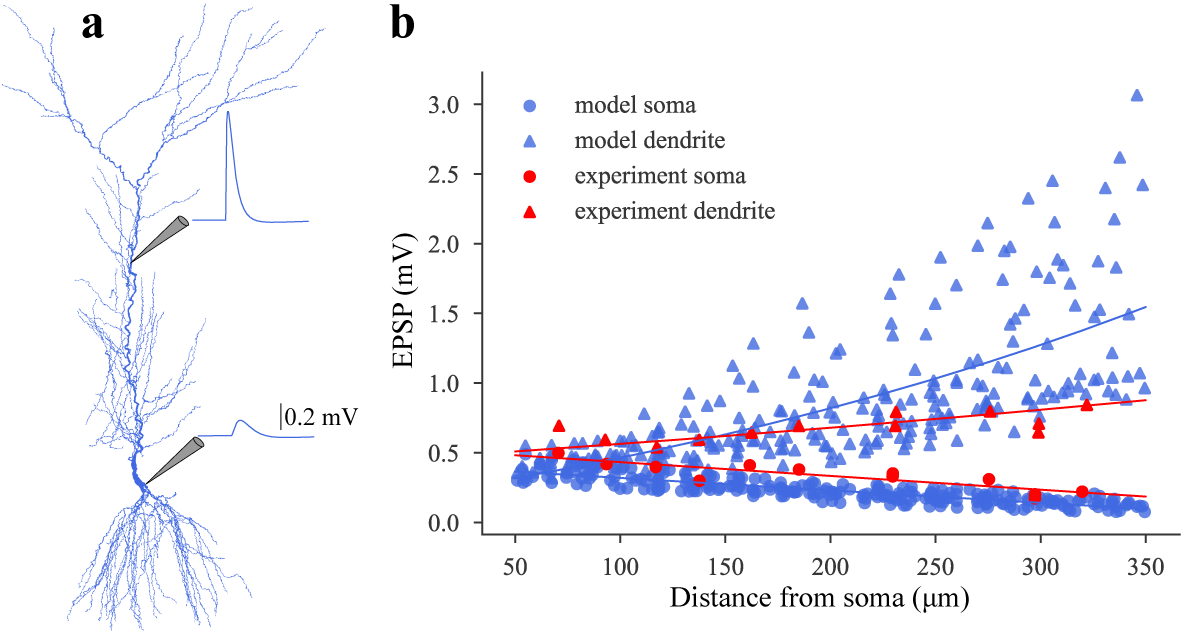
PSP attenuation. Validation of PSP attenuation against experimental data from Magee and Cook (2000). **a:** EPSC like currents were injected to the apical dendrites of the different pyramidal cell models from Migliore et al. (2018) and PSPs were measured at the injection site and at the soma. **b:** Summary of all models injected at different sites (in blue) and comparison to experimental data (in red).

**Figure S2:**
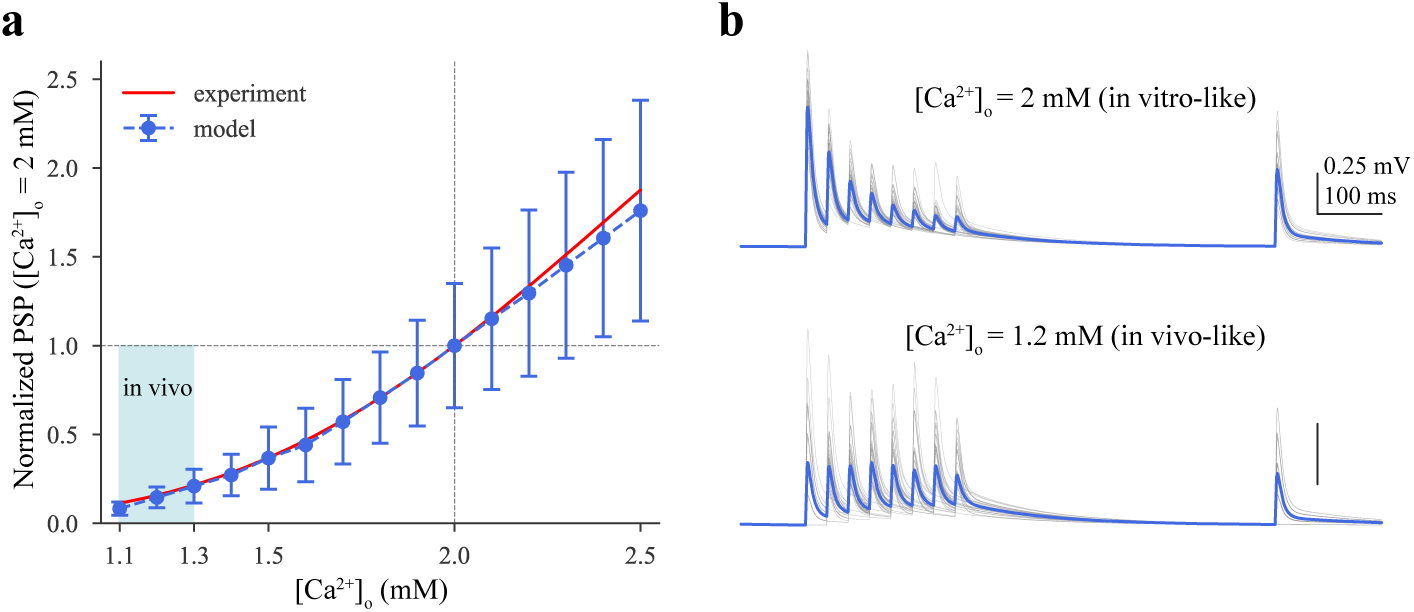
Calcium sensitivity of synaptic physiology. **a:** PC to PC PSP amplitudes at different extracellular calcium concentrations (normalized to 2 mM). Red curve indicates the experimentally measured scaling function which was applied to scale the *U*_*SE*_ parameter of the TM model. Shaded light blue area indicates the *in vivo* range 1.1-1.3 mM. **b:** Same *in silico* PC to PC pair at two different extracellular calcium concentrations. *In vitro* like is shown on top, while the *in vivo* one at the bottom. Single trials (n=35) are shown in gray and their average in blue. Postsynaptic cells were held at −65 mV in *in silico* current-clamp mode.

## Supplementary Tables

**Table S1:** Summary of paired recording experiments from CA1 in voltage-clamp mode (PSCs in pA). Holding potentials are not corrected for the indicated liquid junction potential. * in the rise time constant column indicates 20-80% rise time, instead of 10-90%. Abbreviations are as in **Figure 2 b**

| Presyn. | Postsyn. | Animal | Elect. | Ampl.<br>(pA) | Rise<br>(ms) | Decay<br>(ms) | Hold.<br>(mV) | [Ca<sup>2+</sup>]<br>(mM) | Temp.<br>[°C] | LJP<br>(mV) | Reference |
| --- | --- | --- | --- | --- | --- | --- | --- | --- | --- | --- | --- |
| AA | PC | rat (P10-17) | patch | 308±103 | 0.8±0.1 | 11.2±0.9 | -70 | 3 | ~30 | - | Maccaferri et al. (2000) |
| BS | PC | rat (P10-17) | patch | Fig6) A,B | 2±0.2 | 16.1±1.1 | -70 | 3 | ~30 | - | Maccaferri et al. (2000) |
| CCKBC | PC | rat (P16-20) | patch | 118±13 | 0.73±0.05 | 6.8±0.2 | -70 | 2 | 33 | - | Neu et al. (2007) |
| CCKBC | PC | rat (P16-21) | patch | 53.7±17.2 | - | - | -70 | 2 | 33 | - | Földy et al. (2007) |
| CCKBC | PC | rat (P17-22) | patch | 115.4±10.8 | 0.63±0.04 | 6.47±0.27 | -58.3±0.5 | 2 | 33 | - | Lee et al. (2010) |
| NGF | PC | rat (P18-24) | patch | 4.9±1 | - | 50±4.9 | -50±2 | 2.5 | 33±2 | 12 | Price et al. (2008) |
| OLM | PC | rat (P10-17) | patch | 26±10 | 6.2±0.6 | 20.8±1.7 | -70 | 3 | ~30 | - | Maccaferri et al. (2000) |
| PVBC | PC | rat (P16-21) | patch | 43.6±17.9 | - | - | -70 | 2 | 33 | - | Földy et al. (2007) |
| SCA | PC | rat (P17-22) | patch | 60.2±8.1 | 1.43±0.12 | 8.3±0.44 | -58.1±0.8 | 2 | 33 | - | Lee et al. (2010) |
| NGF | NGF | rat (P12-21) | patch | 20.97±23.68 |  | 42.05±22.03 | -50 | 2.5 | 33-35 | - | Price et al. (2005) |
| NGF | NGF | rat (P18-22) | patch | 85.3±9.6 | 3.69±0.34* | 60.3±4.7 | -65 | 2 | 33±1 | - | Karayannis et al. (2010) |
| OLM | NGF | rat (P18-20) | patch | 19.2 | 2.2 | 10.8 | -50 | 2 | 33±1 | - | Elfant et al. (2008) |
| OLM | PVBC | rat (P18-20) | patch | 11.7±1 | 2.6±1.3 | 16.5±3.9 | -50 | 2 | 33±1 | - | Elfant et al. (2008) |
| OLM | SCA | rat (P18-20) | patch | 19.5±4.7 | 1.9±0.4 | 31.2±4.5 | -50 | 2 | 33±1 | - | Elfant et al. (2008) |

**Table S2:** Summary of paired recording experiments from CA1 in current-clamp mode (PSPs in mV). No LJP correction is necessary since all recordings were obtained with sharp electrodes. * in the half width column indicates decay time constant instead of half width. Abbreviations are as in **Figure 2 b**

| Presyn. | Postsyn. | Animal | Elect. | Ampl.<br>(mV) | Rise<br>(ms) | HalfW<br>(ms) | Hold.<br>(mV) | [Ca<sup>2+</sup>]<br>(mM) | Temp.<br>[°C] | LJP<br>(mV) | Reference |
| --- | --- | --- | --- | --- | --- | --- | --- | --- | --- | --- | --- |
| PC | PC | rat (100-180g) | sharp | 0.7±0.5 | 2.7±0.19 | 16.8±4.1 | -67-70 | 2.5 | 34-36 | Ø | Deuchars and Thomson (1996) |
| AA | PC | rat (90-160g) | sharp | 0.51±0.07 | 5±0.2 | 45.6±2 | -53±1 | 2.5 | 34-36 | Ø | Pawelzik et al. (1999) |
| BS | PC | rat (90-160g) | sharp | 0.86±0.55 | 8.5±3.6 | 43.9±13.9 | -57.6±4.4 | 2.5 | 34-36 | Ø | Pawelzik et al. (1999) |
| BS | PC | rat (120-200g) | sharp | 0.55±0.15 | 7.4±1.4 | 54.6±4.2 | -58.5±0.5 | 2. | 34-36 | Ø | Pawelzik et al. (2002) |
| BS | PC | rat (140-200g) | sharp | 0.8±0.6 | 8.4±3.2 | 42.1±17 | -53 | 2.5 | 33±1 | Ø | Fuentelba et al. (2008) |
| CCKBC | PC | rat (120-200g) | sharp | 1.17±0.44 | 5.4±2.5 | 35.5±19.5 | -65-85 | 2.5 | 34-36 | Ø | Ali et al. (1999) |
| CCKBC | PC | rat (100-200g) | sharp | 1.47±1.06 | 6±2.2 | 47.6±13.3 | -55-60 | 2.5? | 34-36 | Ø | Thomson et al. (2000) |
| CCKBC | PC | rat (120-200g) | sharp | 0.7±0.5 | 6.5±1.5 | 44.2±10.1 | -58.6±3.3 | 2.5 | 34-36 | Ø | Pawelzik et al. (2002) |
| Ivy | PC | rat (140-200g) | sharp | 0.8±0.4 | 2.8±0.2 | 54.1±13.8 | -57 | 2.5 | 33±1 | Ø | Fuentelba et al. (2008) |
| PV+BC | PC | rat (150g<) | sharp | 0.45±0.24 | 4.6±3.2 | 32.4±18* | -57.8±4.6 | 2 | 34-35 | Ø | Buhl et al. (1995) |
| PV+BC | PC | rat (120-200g) | sharp | 1.17±0.57 | 4.5±2 | 30.4±11.6 | -65-85 | 2.5 | 34-36 | Ø | Ali et al. (1999) |
| PV+BC | PC | rat (90-160g) | sharp | 0.81±0.92 | 6.8±2.7 | 47.2±16.9 | -54.7±3.8 | 2.5 | 34-36 | Ø | Pawelzik et al. (1999) |
| PVBC | PC | rat (100-200g) | sharp | 1.12±0.74 | 5.1±1.8 | 39.5±15.2 | -55-60 | 2.5? | 34-36 | Ø | Thomson et al. (2000) |
| PVBC | PC | rat (120-200g) | sharp | 0.83±0.37 | 5.13±2.06 | 38.32±12 | -58.4±3 | 2.5 | 34-36 | Ø | Pawelzik et al. (2002) |
| SCA | PC | rat (120-200g) | sharp | 0.38 | 10±2.8 | 45±2.2 | -58.5±0.5 | 2.5 | 34-36 | Ø | Pawelzik et al. (2002) |
| Tri | PC | rat (120-200g) | sharp | 0.8 | 5.6 | 48.8 | -58.5±0.5 | 2.5 | 34-36 | Ø | Pawelzik et al. (2002) |
| PC | BS | rat (90-180g) | sharp | 3.4±3.1 | 1.2±0.5 | 7.6±2.6 | -66 | 2.5 | 34-35 | Ø | Ali et al. (1998) |
| PC | BS | rat (120-200g) | sharp | 0.95±0.3 | 1.2±0.2 | 10.4±1.6 | -66±1 | 2.5 | 34-36 | Ø | Pawelzik et al. (2002) |
| PC | BS | rat (140-200g) | sharp | 1.8±2.3 | 1.5±0.3 | 6.4±2.7 | -60 | 2.5 | 33±1 | Ø | Fuentelba et al. (2008) |
| PC | CCKBC | rat (120-200g) | sharp | 2±2.1 | 1±0.4 | 6.1±1.5 | -67±3 | 2.5 | 34-36 | Ø | Pawelzik et al. (2002) |
| PC | Ivy | rat (140-200g) | sharp | 2.9±2.2 | 1.5±0.3 | 11.5±1.5 | -60 | 2.5 | 33±1 | Ø | Fuentelba et al. (2008) |
| PC | OLM | rat (90-150g) | sharp | 0.93±1.06 | 1.2±0.5 | 7.5±0.7 | -70±2.3 | 2.5 | 34-36 | Ø | Ali and Thomson (1998) |
| PC | PVBC | rat (90-180g) | sharp | 1.4±1.05 | 0.88±0.44 | 5.4±2.2 | -66 | 2.5 | 34-35 | Ø | Ali et al. (1998) |
| PC | PVBC | rat (120-200) | sharp | 3.51±2.9 | 1±0.3 | 5.74±1.78 | -67 | 2.5 | 34-36 | Ø | Pawelzik et al. (2002) |
| BC | BS | rat (150g) | sharp | 0.37 | 1 | 5.6 | -55 | 2 | 34-35 | Ø | Cobb et al. (1997) |
| BC | BS | rat (120-180g) | sharp | 1±0.4 | 1.65±0.5 | 15.6±2.8 | -63±4.4 | 2.5 | 34-36 | Ø | Pawelzik et al. (2003) |
| BC | BC | rat (150g) | sharp | 0.25 | 1.3 | 27 | -59 | 2 | 34-35 | Ø | Cobb et al. (1997) |
| BC | BC | rat (120-180g) | sharp | 1.1±0.47 | 2.5±0.9 | 18.7±9.1 | -59±4 | 2.5 | 34-36 | Ø | Pawelzik et al. (2003) |
| BS | BC | rat (120-180g) | sharp | 0.7±0.4 | 2.5±0.8 | 19.1±9.5 | -59.7±2.7 | 2.5 | 34-36 | Ø | Pawelzik et al. (2003) |
| SCA | SCA | rat (120-200g) | sharp | 0.5 | 5 | 34.3 | -58 | 2.5 | 34-36 | Ø | Pawelzik et al. (2002) |
| SCA | SCA | rat (P18-23) | patch | 0.6±0.41 | 7.0±1.38 | 41.1±12.5 | -55 | 2 | 20-22 | - | Ali (2007) |

**Table S3:**
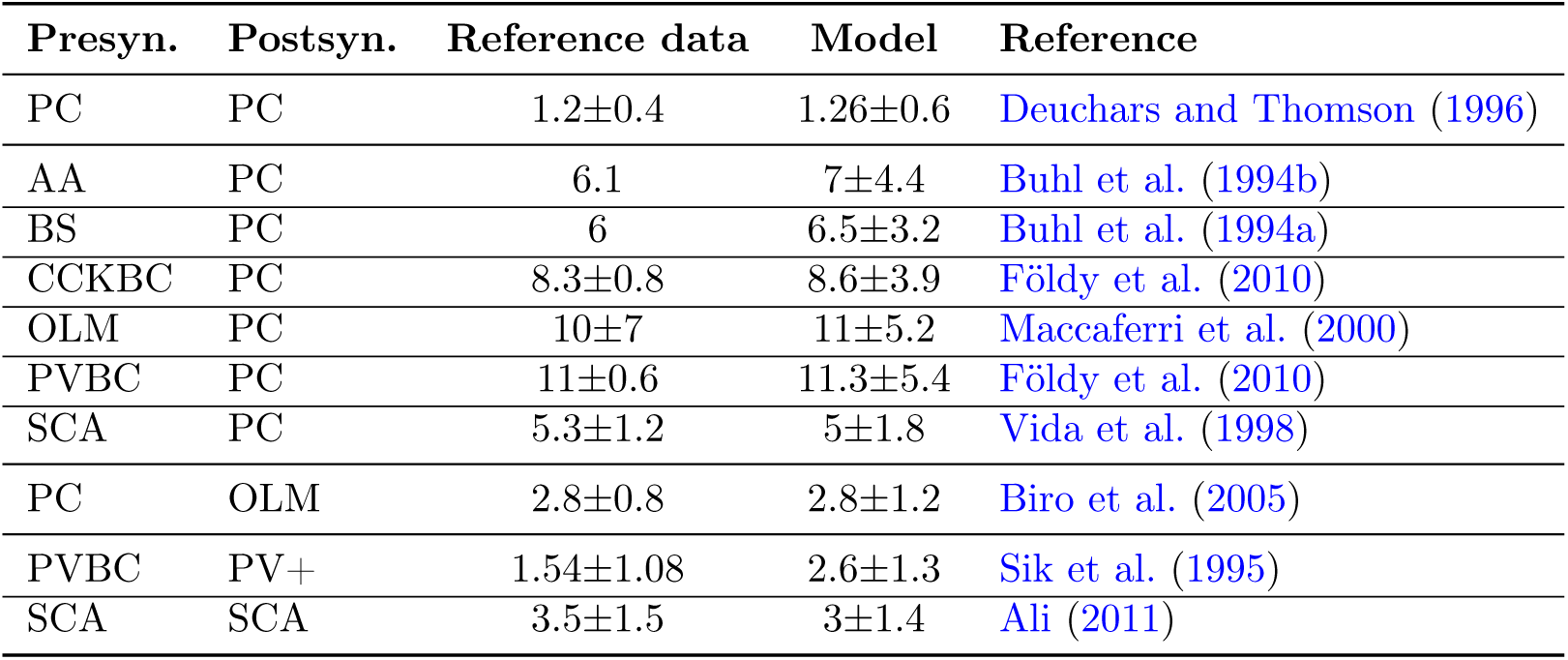
Validation of number of synapses per connections. See **Figure 2 b**). Abbreviations are as in **Figure 2 b**

**Table S4:** Validation of the CV of first PSC amplitudes. See **Figure 3 b**). Abbreviations are as in **Figure 2 b**

| Presyn. | Postsyn. | Reference data | Model | Reference |
| --- | --- | --- | --- | --- |
| AA | PC | $0.29 \pm 0.11$ | $0.28 \pm 0.13$ | Kohus et al. (2016) |
| CCKBC | PC | $0.43 \pm 0.14$ | $0.36 \pm 0.1$ | Kohus et al. (2016) |
| PVBC | PC | $0.26 \pm 0.06$ | $0.28 \pm 0.07$ | Kohus et al. (2016) |
| SCA | PC | $0.38 \pm 0.11$ | $0.31 \pm 0.08$ | Kohus et al. (2016) |
| CCKBC | CCKBC | $0.18 \pm 0.16$ | $0.18 \pm 0.1$ | Kohus et al. (2016) |
| PVBC | AA | $0.45 \pm 0.11$ | $0.17 \pm 0.09$ | Kohus et al. (2016) |
| PVBC | PVBC | $0.17 \pm 0.05$ | $0.22 \pm 0.02$ | Kohus et al. (2016) |

**Table S5:** Validation of PSP amplitudes. See **Figure 3 d**). PC to CCKBC and PC to Ivy are not shown on the figure for visualization purpose. In some cases (indicated with *) outliers were removed from the reference data (see published reference data in Table S1). Abbreviations are as in **Figure 2 b**

| Presyn. | Postsyn. | Reference data (mV) | Model (mV) | Reference |
| --- | --- | --- | --- | --- |
| PC | PC | $0.7 \pm 0.5$ | $0.78 \pm 0.71$ | Deuchars and Thomson (1996) |
| AA | PC | $0.51 \pm 0.07$ | $0.55 \pm 0.17$ | Pawelzik et al. (1999) |
| BS | PC | $0.55 \pm 0.15$ | $0.57 \pm 0.21$ | Pawelzik et al. (2002) |
| CCKBC | PC | $0.7 \pm 0.5$ | $0.67 \pm 0.3$ | Pawelzik et al. (2002) |
| Ivy | PC | $0.8 \pm 0.4$ | $0.8 \pm 0.27$ | Fuentelba et al. (2008) |
| PVBC | PC | $0.83 \pm 0.37$ | $0.76 \pm 0.26$ | Pawelzik et al. (2002) |
| SCA | PC | 0.38 | $0.43 \pm 0.18$ | Pawelzik et al. (2002) |
| Tri | PC | 0.8 | $0.8 \pm 0.36$ | Pawelzik et al. (2002) |
| PC | BS | $0.95 \pm 0.3$ | $1 \pm 0.54$ | Pawelzik et al. (2002) |
| PC | CCKBC | $2 \pm 2.1$ | $1.9 \pm 1.35$ | Pawelzik et al. (2002) |
| PC | Ivy | $2.9 \pm 2.2$ | $2.45 \pm 1.5$ | Fuentelba et al. (2008) |
| PC | OLM | $0.3 \pm 0.13^*$ | $0.25 \pm 0.16$ | Ali and Thomson (1998) |
| PC | PVBC | $1 \pm 0.4^*$ | $1 \pm 0.7$ | Ali et al. (1998) |
| (PV)BC | (PV)BC | 0.25 | $0.25 \pm 0.12$ | Cobb et al. (1997) |

